# Staphylococcal protein A inhibits IgG-mediated phagocytosis by blocking the interaction of IgGs with FcγRs and FcRn

**DOI:** 10.1101/2022.01.21.477287

**Authors:** Ana Rita Cruz, Arthur E. H. Bentlage, Robin Blonk, Carla J. C. de Haas, Piet C. Aerts, Lisette M. Scheepmaker, Inge G. Bouwmeester, Anja Lux, Jos A. G. van Strijp, Falk Nimmerjahn, Kok P. M. van Kessel, Gestur Vidarsson, Suzan H. M. Rooijakkers

## Abstract

Immunoglobulin G molecules are crucial for the human immune response against bacterial infections. IgGs can trigger phagocytosis by innate immune cells, like neutrophils. To do so, IgGs should bind to the bacterial surface via their variable Fab regions and interact with Fcγ receptors (FcγRs) and complement C1 via the constant Fc domain. C1 binding to IgG-labeled bacteria activates the complement cascade, which results in bacterial decoration with C3-derived molecules that are recognized by complement receptors (CRs) on neutrophils. Next to FcγRs and CRs on the membrane, neutrophils also express the intracellular neonatal Fc receptor (FcRn). We previously reported that staphylococcal protein A (SpA), a key immune evasion protein of *Staphylococcus aureus*, potently blocks IgG-mediated complement activation and killing of *S. aureus* by interfering with IgG hexamer formation. SpA is also known to block IgG-mediated phagocytosis in absence of complement but the mechanism behind it remains unclear. Here we demonstrate that SpA blocks IgG-mediated phagocytosis and killing of *S. aureus* through inhibition of the interaction of IgGs with FcγRs (FcγRIIa and FcγRIIIb, but not FcγRI) and FcRn. Furthermore, our data show that multiple SpA domains are needed to effectively block IgG1-mediated phagocytosis. This provides a rationale for the fact that SpA from *S. aureus* contains four to five repeats. Taken together, our study elucidates the molecular mechanism by which SpA blocks IgG-mediated phagocytosis and supports the idea that next to FcγRs, also the intracellular FcRn receptor is essential for efficient phagocytosis and killing of bacteria by neutrophils.

## Introduction

Immunoglobulin G (IgG) antibodies play a key role in the host immune response against bacteria. IgGs consist of two functional domains: the antibody-binding fragment (Fab) and the crystallizable fragment (Fc). Via their variable Fab domain, antibodies can directly neutralize the function of bacterial virulence factors. Moreover, when antibodies bind to the bacterial surface via their Fab domain, their constant Fc domain can trigger bacterial clearance by interacting with the innate immune system (1). IgG-Fcs have two important effector functions. While they can directly bind to Fc gamma receptors (FcγRs) expressed on the surface of innate immune cells, they can also bind complement C1 via clustered IgGs and activate the classical complement pathway. The activation of the complement cascade results in bacterial decoration with C3-derived opsonins, that are in turn recognized by complement receptors (CRs) on innate immune cells, like neutrophils. Both pathways ultimately trigger phagocytosis and killing of the invading bacteria.

FcγRs are membrane glycoproteins and are divided into six classes: FcγRI (CD64), FcγRIIa (CD32a), FcγRIIb (CD32b), FcγRIIc (CD32c), FcγRIIIa (CD16a), and FcγRIIIb (CD16b). FcγRI is the only high-affinity receptor, as it can bind to monomeric IgGs while the other receptors mainly bind to aggregated IgGs. The low affinity receptors have polymorphic variants. Besides the classical extracellular Fc receptors, IgG-Fcs are also recognized by the intracellular neonatal Fc receptor (FcRn) (2) (see **Fig. 1A**). FcRn is found on different cell types, including epithelial cells, endothelial cells and placental syncytiotrophoblasts. FcRn is mainly known for its role in transferring IgG from the mother to the fetus (3) and in the regulation of IgG half-life (4, 5). More recently, FcRn was also found to be expressed in monocytes, macrophages, dendritic cells (6) and neutrophils (2) and shown to be involved in phagocytosis of IgG-coated pneumococci (2). The binding sites of FcγRs and FcRn on IgG are different (see **Fig. 1B**): while FcγRs bind with a 1:1 stoichiometry to the lower hinge and CH2 domain of IgG (7), FcRn binds the CH2-CH3 interface of IgG with a 2:1 stoichiometry (8–10). Another difference is that FcRn-IgG binding only occurs at acidic pH (< 6.5) (11).

**Figure 1.**
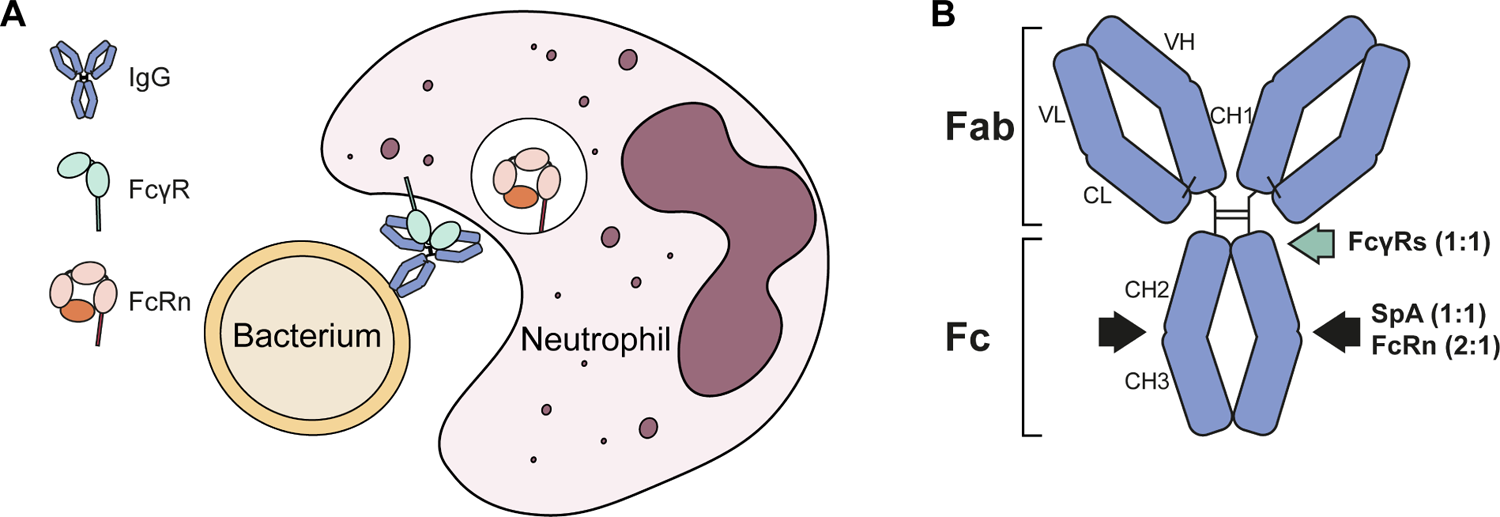
Neutrophils express FcγRs and FcRn that recognize IgG-Fcs in structurally distant sites. (A) Schematic representation of an IgG-labeled bacterium being phagocytized by a neutrophil, showing binding of extracellular FcγRs to IgGs and FcRn inside granular structures. (B) Schematic illustration of IgG indicating the binding regions of staphylococcal protein A (SpA), FcγRs and FcRn. FcγRs bind to the lower hinge and CH2 domain of IgG with a 1:1 stoichiometry while SpA and FcRn bind to the CH2-CH3 interface with a 1:1 and 2:1 stoichiometry, respectively.

Interestingly, the Fc fragment of IgGs is not only recognized by host immune effector proteins, but also by bacterial immune evasion molecules (12), like staphylococcal protein A (SpA). SpA is a key immune evasion factor of *Staphylococcus aureus*, a prominent human pathogen that spreads in healthcare facilities and in the community, causing multiple diseases (13).

SpA is mainly anchored to the bacterial cell wall, although it is also found in the extracellular milieu (14, 15). SpA is composed of five highly homologous three-helix-bundle domains (named A to E), each of which can bind the CH2-CH3 interface of IgG (see **Fig. 1B** and **Fig. 2A**), via helices I and II (16). Moreover, SpA domains also bind the Fab region of most VH3-type family of antibodies, via helices II and III (17). Of note, the Fc domain recognition properties of SpA are subclass and allotype specific. SpA binds IgG1, IgG2 and IgG4 subclasses, but not to the majority of IgG3 allotypes (18). This is due to an amino acid substitution in position 435, where an histidine in IgG1, IgG2 and IgG4 becomes an arginine in most of IgG3 allotypes (19, 20). Therefore, the effector functions of IgG3 remain unaffected by the presence of SpA (21).

**Figure 2.**
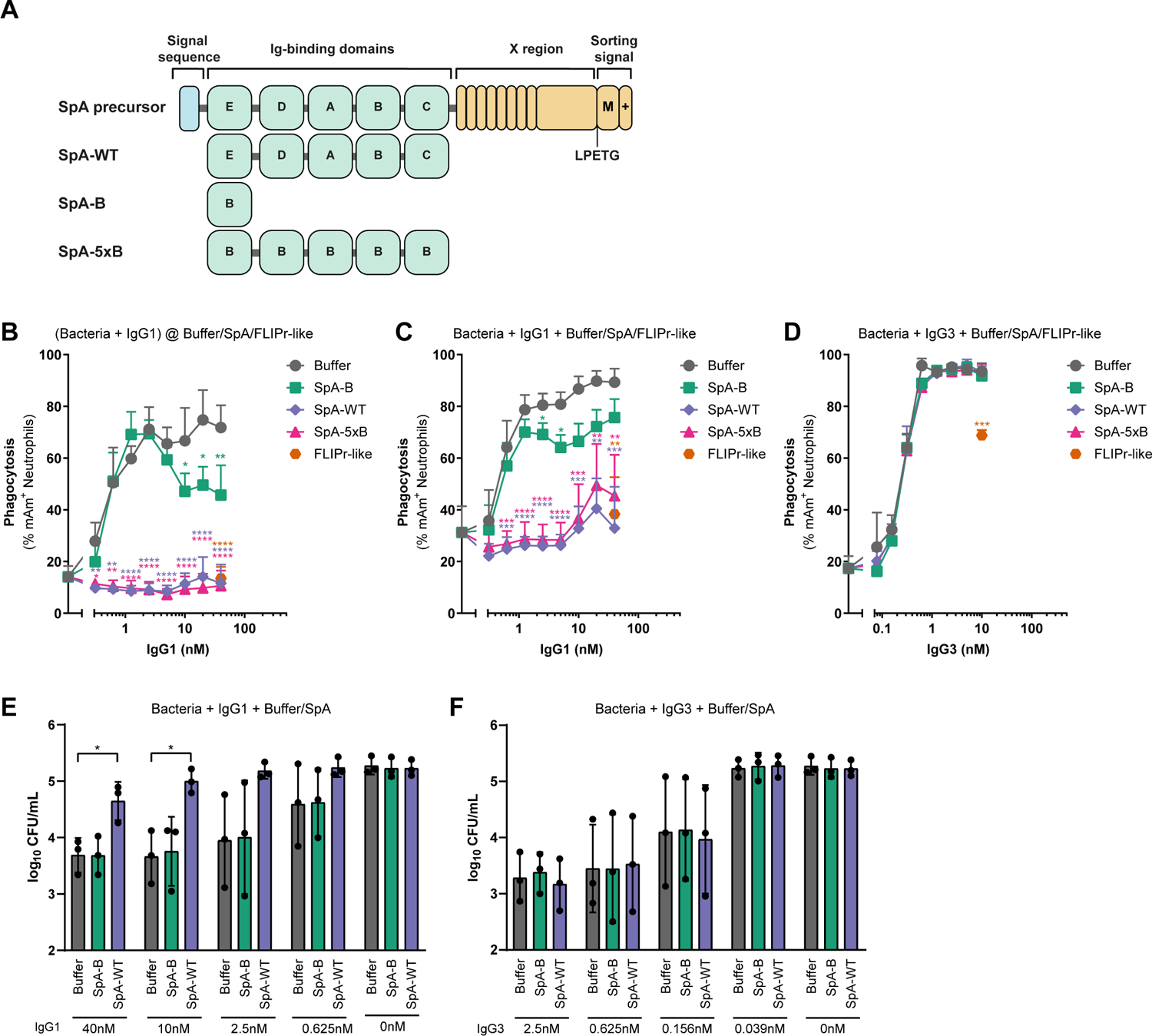
Soluble SpA requires multiple domains to effectively block IgG1-mediated phagocytosis and killing of S. aureus. (A) Schematic representation of SpA precursor and recombinant SpA proteins used in this study. Unprocessed SpA consists of a signal sequence, five Ig-binding domains (E, D, A, B, and C), an X region and a sorting region. The recombinant SpA proteins used here include solely the Ig-binding domains. While SpA-WT is composed of five different Ig-binding domains, SpA-B contains a single B domain and SpA-5xB consists of five repeating B domains. (B) Phagocytosis of *S. aureus* Newman Δ*spa*/*sbi* after incubation of bacteria with a concentration range of anti-WTA IgG1, followed by buffer (grey), 200 nM of SpA-B (green), SpA-WT (blue), SpA-5xB (pink) or FLIPr-like (orange), measured by flow-cytometry. Bacteria were washed after incubation with IgGs to remove unbound IgG and only after buffer, SpA or FLIPr-like was added. (C, D) Phagocytosis of *S. aureus* Newman Δ*spa*/*sbi* after incubation of bacteria with a concentration range of anti-WTA IgG1 (C) or IgG3 (D), in absence (buffer; grey) or presence of 200 nM soluble SpA-B (green), SpA-WT (blue), SpA-5xB (pink) or FLIPr-like (orange), measured by flow-cytometry. Bacteria, IgG and buffer, SpA or FLIPr-like were incubated at the same step. (E, F) CFU enumeration of Newman Δ*spa*/*sbi* after incubation with anti-WTA IgG1 (E) or IgG3 (F) in absence (buffer; grey) or presence of 200 nM SpA-B (green) or SpA-WT (blue), followed by incubation with human neutrophils. Bacteria, IgG and buffer, SpA or FLIPr-like were incubated at the same step. Data are presented as % of mAm^+^ PMNs ± SD of three (B, C) or two (D) independent experiments, or as log_10_ CFU/mL ± SD of three independent experiments (E, F). Statistical analysis was performed using one-way ANOVA to compare buffer condition with SpA-B, SpA-WT, SpA-5xB and FLIPr-like conditions and displayed only when significant as *P ≤ 0.05; **P ≤ 0.01; ***P ≤ 0.001; ****P ≤ 0.0001.

We and others showed that SpA protects *S. aureus* from phagocytic killing by binding the IgG-Fc fragment (21–23). In our previous study, we unveiled how SpA blocks IgG-mediated complement activation and subsequent killing of *S. aureus* (21). We showed that SpA binds competitively to the Fc-Fc interaction interface on IgG monomers, which effectively prevents IgGs from forming IgG hexamers (21). IgG hexamerization on antigenic surfaces is important for efficient binding of C1 and subsequent activation of the complement system (24).

However, SpA was also reported to prevent IgG-dependent phagocytic killing in the absence of complement (23). To date, the precise mechanism by which SpA blocks FcγR-mediated phagocytosis remains elusive. SpA is known to reduce binding of antigen-complexed and heat-aggregated IgG to Fc receptor-bearing cells (25). However, soluble FcγRI and FcγRIIa were shown not to compete with SpA for binding to soluble IgG (26). Here, we investigate how SpA interferes with FcγR-mediated phagocytosis of *S. aureus* and reveal an important role of FcRn in this process.

## Materials and Methods

### Production of human monoclonal antibodies

The human anti-DNP and anti-WTA GlcNAc-β-4497 monoclonal antibodies were produced recombinantly in human Expi293F cells (Life Technologies) as described before (21). The VH and VL sequences of the anti-DNP (DNP-G2a2) (27) and anti-WTA GlcNAc-β-4497 (patent WO/2014/193722) (28) were derived from previously reported antibodies. The anti-Hla (MEDI4893; patent WO/2017/075188A2), which served as a control, was a gift from Dr Alexey Ruzin, MedImmune (AstraZeneca). The human anti-TNFα IgG1 monoclonal antibody was expressed in HEK-293F FreeStyle cell line expression system (Life Technologies, Carlsbad, CA) with co-transfection of vectors encoding p21, p27, and pSVLT genes to increase protein production (29). The VH and VL sequences of the anti-TNFα (patent US 6090382A) were also derived from a previously published antibody. Anti-TNFα IgG1 was purified on Protein A HiTrapHP columns (GE Healthcare Life Sciences, Little Chalfront, UK) using Akta-prime plus (GE Healthcare Life Sciences) and dialyzed overnight against PBS.

### Cloning, expression and purification of staphylococcal proteins

SpA constructs were cloned, expressed and purified as reported before (21). The wild-type B domain of SpA (SpA-B), SpA-B lacking Fc-binding properties (SpA-B^KK^; Q9K and Q10K mutations) and SpA-B lacking Fab-binding properties (SpA-B^AA^; D36A and D37A mutations) were produced for a previous study (21), while the five-domains SpA (SpA-WT) and the repeating five B domains SpA (SpA-5xB) were newly produced, following the same steps described before (21). For the design of SpA-5xB, multiple attempts were made to rearrange its nucleotide sequence to make the synthesis as the gBlock (Integrated DNA technologies, IDT) possible. Besides the in house produced SpA-WT, for some of the experiments, we used a five-domains SpA (also named SpA-WT) that is commercially available (Prospec, PRO-1925). The recombinant protein FLIPr-like was expressed and purified as described before (30).

### Bacterial strains and culture conditions

mAmetrine (mAm)-labeled *S. aureus* Newman Δ*spa*/*sbi* and Newman WT were constructed as described before (32). Briefly, bacteria were transformed with a pCM29 plasmid that constitutively and robustly expresses a codon optimized mAm protein (GenBank: KX759016) (33) from the sarAP1 promoter (34). For generation of p*spa* complemented Newman Δ*spa*/*sbi*, the *spa* gene and its promoter were first PCR amplified from genomic DNA of *S. aureus* Newman WT. The PCR products were cloned into the pCM29 vector via Gibson assembly and *E. coli* DC10b transformed with pCM29-*spa* by heat shock. Subsequently, the plasmid was isolated and competent *S. aureus* Newman Δ*spa/sbi* were transformed with the plasmid through electroporation using Bio-Rad Gene Pulser Xcell Electroporation System (200 Ω, 25 µF, 2.5 kV). After recovery, bacteria were plated on Todd-Hewitt agar supplemented with 5 μg/mL chloramphenicol to select plasmid-complemented colonies. Bacteria were grown overnight in Todd Hewitt broth (THB) plus 10 μg/mL chloramphenicol, diluted to an OD_600_ = 0.05 in fresh THB plus chloramphenicol, and cultured until midlog phase (OD_600_ = 0.5). The mAmetrine-expressing strains Newman Δ*spa*/*sbi* and Newman WT were washed and resuspended in RPMI-H medium (RPMI + 0.05% HSA) and stored until use at −20 °C. The Newman *Δspa/sbi* + p*spa* strain was FITC-labeled before storage. Briefly, midlog phase bacteria were washed with PBS and resuspended in 0.5 mg/ml FITC (Sigma Aldrich) in PBS for 1 h on ice, washed twice in PBS, resuspended in RPMI-H medium and stored until use at −20 °C.

### Phagocytosis of *S. aureus* by neutrophils

Human neutrophils were purified from blood of healthy donors by the Ficoll/Histopaque density gradient method (35). To study the inhibitory effect of SpA on phagocytosis, we used a recently described phagocytosis assay (32), with some adaptations. mAm-expressing Newman Δ*spa*/*sbi* (7.5 × 10^5^ CFU) were first incubated with human monoclonal anti-WTA IgG1 for 15 min at 37 °C with shaking (±700 rpm) in a round-bottom microplate. After, bacteria were washed with RPMI-H by centrifugation (3600 rpm, 7 min) and incubated in absence or presence of SpA-B, SpA-WT, SpA-5xB or FLIPr-like for 15 min at 37 °C with shaking. Finally, bacteria were mixed with neutrophils for another 15 min at 37 °C with shaking, at a 10:1 bacteria:neutrophil ratio. Alternatively, bacteria were simultaneously incubated with human monoclonal anti-WTA IgG1, IgG3 or heat-inactivated normal human serum (HI-NHS) with buffer, SpA constructs or FLIPr-like in RPMI-H medium for 15 min at 37 °C with shaking. Bacteria were then incubated with freshly isolated neutrophils for another 15 min at 37 °C with shaking. To evaluate the inhibitory effect of cell-attached SpA on phagocytosis, 7.5 × 10^5^ CFU of fluorescently labeled Newman Δ*spa*/*sbi*, Newman WT or Newman Δ*spa*/*sbi* + p*spa* were incubated with anti-WTA IgG1, IgG3 or HI-NHS in RPMI-H. After 15 min at 37 °C with shaking, IgG-opsonized bacteria were incubated with 7.5 × 10^4^ neutrophils for another 15 min at 37 °C with shaking. All samples were fixed with 1% paraformaldehyde in RPMI-H (final concentration). The binding/internalization of mAm-bacteria to the neutrophils was detected using flow cytometry (BD FACSVerse) and data were analyzed based on FSC/SSC gating of neutrophils using FlowJo software.

### Killing of *S. aureus* by neutrophils

mAmetrine-expressing Newman Δ*spa*/*sbi* were freshly grown to midlog phase, washed with PBS and resuspended in HBSS-H medium (Hank’s balanced salt solution (HBSS) + 0.1% HSA). Newman Δ*spa*/*sbi* (8.5 × 10^5^ CFU) were incubated with fourfold titration of anti-WTA IgG1 or IgG3 in the absence or presence of 200 nM SpA-B or SpA-WT in HBSS-H. After 30 min at 37 °C, bacteria were incubated with neutrophils for 90 min under 5% CO2 at 37 °C, at a 1:1 bacteria:neutrophil ratio. Subsequently, neutrophils were lysed with cold 0.3% (wt/vol) saponin in water for up to 15 min on ice. Samples were serially diluted in PBS and plated in duplicate onto TSA plates, which were incubated overnight at 37 °C. Viable bacteria were quantified by CFU enumeration.

### ELISA assays

MaxiSorp plates (Nunc) were coated with 3 µg/mL SpA-B, SpA-B^KK^, SpA-B^AA^, SpA-WT or SpA-5xB in 0.1 M sodium carbonate at 4 °C, overnight. After three washes with PBS-T (PBS, 0.05% (v/v) Tween-20) pH 7, the wells were blocked with 4% bovine serum albumin (BSA) in PBS-T, for 1 h at 37 °C. The following incubations were performed for 1 h at 37 °C followed by three washes with PBS-T. 1 µg/mL of anti-WTA IgG1, anti-WTA IgG3, or anti-Hla IgG1 were diluted in 1% BSA in PBS-T and added to the wells. For IgG-SpA binding detection at neutral and acidic conditions, the wells were incubated with a concentration range of a-WTA IgG1 (5-fold serial dilutions starting from 20 nM) diluted in PBS-T at pH 7 or pH 6. Bound antibodies were detected with horseradish peroxidase (HRP)-conjugated goat F(ab’)₂ anti-human kappa (Southern Biotech) in 1% BSA in PBS-T and Tetramethylbenzidine as substrate. The reaction was stopped with 1N sulfuric acid and absorbance was measured at 450 nm in the iMarkTM Microplate Absorbance Reader (BioRad).

### SpA expression and antibody binding on *S. aureus* Newman strains

To detect cell-attached SpA, 7.5 × 10^5^ CFU of fluorescently labeled *S. aureus* Newman strains (Δ*spa*/*sbi*, WT and Δ*spa*/*sbi* + p*spa*) were incubated in a round-bottom microplate with 1 µg/mL biotin-conjugated chicken anti-Protein A (Immunology Consultants Laboratory) in RPMI-H for 30 min at 4 °C under shaking conditions (±700 rpm), washed by centrifugation with RPMI-H (3600 rpm, 7 min), and incubated with 2 µg/mL Alexa Fluor^647^-conjugated streptavidin (Jackson ImmunoResearch) in RPMI-H for another 30 min at 4 °C under shaking conditions. To measure antibody binding to the same strains, bacteria (7.5 × 10^5^ CFU) were incubated with 3-fold serial dilutions of anti-DNP IgG1 or IgG3, anti-WTA IgG1 or IgG3 in RPMI-H (starting from 10 nM IgG), for 30 min at 4 °C, shaking. Subsequently, bacteria were washed by centrifugation with RPMI-H (3600 rpm, 7 min), and incubated with 0.5 µg/mL Alexa Fluor^647^-conjugated goat F(ab’)₂ anti-human kappa (Southern Biotech) in RPMI-H for 30 min at 4 °C under shaking conditions. After an additional wash with RPMI-H, all samples were fixed with 1% paraformaldehyde in RPMI-H. SpA expression and antibody binding on *S. aureus* Newman strains were detected using flow cytometry (BD FACSVerse) and data were analyzed using FlowJo software.

### Surface plasmon resonance measurements

Affinity measurements of soluble IgG to FcγRs and FcRn were performed with the IBIS MX96 biosensor system as described previously (36). In short, C-terminally site-specifically BirA-biotinylated human FcγRIIa H131, FcγRIIa R131, FcγRIIb, FcγRIIIa F158, FcγRIIIa V158, FcRn (SinoBiologicals; 10374-H27H1-B, 10374-H27H-B, 10259-H27H-B, 10389-H27H-B, 10389-H27H1-B, CT071-H27H-B, respectively), FcγRIIIb NA1 and FcγRIIIb NA2 (produced at the laboratory of Sanquin) were spotted onto a SensEye G-Streptavidin sensor (Senss, Enschede, Netherlands) using a continuous flow micro spotter (Wasatch Microfluidics, Salt Lake City, UT, United States) in running buffer (PBS 0.0075% Tween-80 (Amresco)), pH 7.4. The receptors were added at the following concentrations: 10 nM of FcγRIIa H131, FcγRIIa R131 and FcγRIIb, 30 nM of FcγRIIIa 158F, FcγRIIIb NA1 and FcγRIIIb NA2 and 100 nM of FcγRIIIa 158V. For IgG-FcγRI binding measurements, 30 nM of biotinylated mouse IgG1 anti-His was first spotted on the sensor, followed by 50 nM of His-tagged FcγRI (SinoBiologicals, 10256-H27H). Anti-TNFα IgG1 (200 nM) was then injected in combination with 1µM or 200 nM SpA-B, SpA-WT, SpA-5xB or running buffer. For IgG-FcRn binding measurements, the running buffer was at pH6. Regeneration after each sample was carried out with 10 mM Gly– HCl, pH 2.4.

### Binding assays with hFcγR-expressing CHO cell lines

CHO cells expressing human FcγRI, FcγRIIa H131, FcγRIIa R131, FcγRIIIb NA1 or FcγRIIIb NA2 were generated at University of Erlangen-Nürnberg laboratory (37). Untransfected CHO cells were maintained in RPMI medium supplemented with 10% FCS, 2 mM glutamine, 1 mM sodium pyruvate, 0.1 mg/mL pen/strep and 0.1 mM non-essential amino acids at 37%, 5% CO2, while for the CHO cell lines stably transfected with human FcγRs, 0.05 mg/mL G418 was also added to the supplemented RPMI medium. Cells were collected by brief trypsinization, washed and resuspended in RPMI-H. Viability was >95% as assessed with trypan blue. mAm-expressing Newman Δ*spa*/*sbi* (7.5×10^5^ CFU or 2.25×10^6^ CFU) were first labeled with anti-WTA IgG1 or buffer control for 30 min at 4 °C with shaking (±700 rpm) in a round-bottom microplate. Bacteria were washed by centrifugation with RPMI-H (3600 rpm, 7 min) and after mixed with buffer, SpA-B, SpA-WT, SpA-5xB or FLIPr-like in RPMI-H for 30 min at 37 °C with shaking. Finally, CHO cells (7.5×10^4^ cells) were added to the mixture (10:1 bacteria:cells ratio for FcγRI-, FcγRIIa H131- and FcγRIIa R131-expressing CHO cells and 30:1 bacteria:cells ratio for FcγRIIIb NA1- and FcγRIIIb NA2-expressing CHO cells) for another 30 min at 37 °C with shaking. The reaction was stopped and fixed by addition of 1% paraformaldehyde (final concentration) in RPMI-H. The binding of mAm-bacteria to the CHO cells was measured by flow cytometry (BD FACSVerse) and data were analyzed based on FSC/SSC gating of CHO cells using FlowJo software.

### FcRn receptor-coated beads assays

Streptavidin beads (Dynabeads M-270; Invitrogen) were washed in PBS-TH (phosphate-buffered saline [PBS], 0.05% [vol/vol] Tween-20, and 0.5% human serum albumin [HSA]) and incubated (diluted 100×) with 1 μg/mL C-terminally site-specifically BirA-biotinylated human FcRn (Acrobiosystems, FCM-H82W7) in PBS-TH for 30 min at 4 °C with shaking. FcRn-labeled beads were then washed twice with PBS-TH and resuspended in PBS-TH at pH 6.0. For binding of bacteria-bound IgG to FcRn-coated beads, mAm-expressing Newman Δ*spa*/*sbi* (6×10^5^ CFU) were first incubated with anti-WTA IgG1 in RPMI-H for 30 min at 4 °C with shaking (±700 rpm). After a single wash by centrifugation (3600 rpm, 7 min) with PBS-TH at pH6, IgG1-labeled bacteria were incubated in absence or presence of SpA-B, SpA-WT or SpA-5xB in PBS-TH at pH6 for 30 min at 37 °C with shaking. From this step, all incubations and washes were performed using PBS-TH at pH6. FcRn-coated beads were then mixed with the bacteria for 30 min at 37 °C with shaking. For each condition, 0.5 μL of FcRn-coated beads were used (∼3 × 10^5^ beads/condition). After two washes with PBS-TH, the beads were incubated with 1 µg/mL Alexa Fluor^647^-conjugated goat F(ab’)_2_ anti-human kappa (Southern Biotech, 2062-31) in PBS-TH for 30 min at 4 °C with shaking. For binding of soluble IgG to FcRn-coated beads, anti-DNP IgG1 was first incubated in absence or presence of recombinant SpA-B, SpA-WT, SpA-5xB or FLIPr-like in a round-bottom microplate at 4 °C with shaking.

After 30 min of incubation, IgG1 + Buffer/ SpA/ FLIPr-like were mixed with FcRn-labelled beads for another 30 min at 37 °C, shaking. After two washes with PBS-TH, the beads were incubated with 1 µg/mL Alexa Fluor^647^-conjugated goat F(ab’)₂ anti-human kappa (Southern Biotech, 2062-31) in PBS-TH for 30 min at 4 °C with shaking. Finally, the beads were washed twice with PBS-TH and then fixed with 1% paraformaldehyde in PBS-TH. Binding of bacteria-bound IgG and soluble IgG to the beads was detected using flow cytometry (BD FACSVerse) and data were analyzed based on single bead population using FlowJo software.

## Ethical Statement

Human serum and blood were obtained from healthy donors after informed consent was obtained from all subjects, in accordance with the Declaration of Helsinki. Approval from the Medical Ethics Committee of the University Medical Center Utrecht was obtained (METC protocol 07-125/C, approved March 1, 2010).

## Statistical analysis

Statistical analysis was performed with GraphPad Prism v.8.3 software, using one-way ANOVA as indicated in the figure legends. At least three experimental replicates were performed to allow statistical analysis.

## Results

### Soluble SpA requires multiple domains to effectively block IgG1-mediated phagocytosis and killing of *S. aureus*

To investigate how SpA blocks IgG-mediated phagocytosis, we first studied whether different forms of soluble SpA affect phagocytosis of *S. aureus* by human neutrophils. To exclusively determine the effect of soluble SpA in this assay, we used an isogenic mutant of Newman *S. aureus* strain lacking both SpA and the second Ig-binding protein of *S. aureus,* Sbi (38) (Newman Δ*spa*/*sbi*). Newman Δ*spa*/*sbi* was first incubated with a human monoclonal IgG1 antibody that targets wall teichoic acid (anti-WTA IgG1). WTA is a highly abundant surface glycopolymer anchored to the peptidoglycan layer of *S. aureus* (39). After a wash to remove unbound IgGs, IgG1-labeled bacteria were incubated with different recombinant SpA constructs: wild-type SpA composed of five IgG-binding domains (SpA-WT), a single SpA-B domain (SpA-B) and a SpA variant composed of five repeating B domains (SpA-5xB) (see **Fig. 2A**). As a positive control we used the homologue of formyl peptide receptor-like 1 inhibitor (FLIPr-like), a staphylococcal phagocytosis inhibitor that directly binds to FcγRs (40). While SpA-WT and SpA-5xB potently reduced phagocytosis mediated by IgG1, the single SpA-B domain showed a minimal effect on phagocytosis (**Fig. 2B** and **S1A**).

To understand whether the SpA constructs could also inhibit phagocytosis when in presence of soluble IgGs, we also assessed phagocytosis when bacteria, IgGs and SpA were incubated at the same step. In this set-up where SpA can interact with both target-bound and soluble antibodies, we observed that multi-domain SpA can still inhibit IgG1-mediated phagocytosis, although the inhibitory effect was slightly weaker (**Fig. 2C** and **S1B**). When instead of anti-WTA IgG1 antibodies we used IgG3, none of the SpA constructs affected phagocytosis (**Fig. 2D**), suggesting that SpA interferes with IgG-mediated phagocytosis by binding to the Fc region of IgGs. Although anti-WTA antibody clone 4497 used in these assays belongs to VH3-type family (41), it does not bind SpA via its Fab region (21). The SpA binding properties of anti-WTA IgG1 and IgG3 were verified by comparing their binding to the wild-type SpA-B with two SpA-B variants that cannot interact with Fc (SpA-B^KK^) or Fab (SpA-B^AA^) domains of IgG (**Fig. S1C, D**). The binding of a VH3 family antibody (anti-Hla IgG1) was also measured as a control for Fab binding to SpA-B^KK^. This confirms that soluble SpA blocks IgG-mediated phagocytosis by binding to the IgG-Fc region.

Next, we evaluated whether inhibition of IgG-mediated phagocytosis by SpA results in less phagocytic killing of *S. aureus* by human neutrophils. Upon engulfment, neutrophils can kill bacteria intracellularly by exposing them to antimicrobial peptides, enzymes and reactive oxygen species (42). While anti-WTA IgG1 antibodies alone induced killing of *S. aureus*, the presence of SpA-WT blocked killing (**Fig. 2E**). In line with the phagocytosis data, we observed that a single SpA-B domain cannot block IgG1-mediated killing (**Fig. 2E**). As expected, the presence of SpA proteins did not affect IgG3-mediated killing of *S. aureus* (**Fig. 2F**).

Altogether, these data show that soluble SpA blocks phagocytosis and killing of *S. aureus* by binding to the Fc region of IgG. Furthermore, we find that multiple SpA domains are required to potently block IgG-mediated phagocytosis.

### Surface-bound SpA blocks IgG-mediated phagocytosis of *S. aureus*

Next, we assessed whether surface attached SpA also reduces IgG-mediated phagocytosis. To do so, we compared phagocytosis of wild-type *S. aureus* strain Newman with Newman Δ*spa*/*sbi*. In addition, we complemented Newman Δ*spa*/*sbi* with SpA, by overexpressing the *spa* gene from a plasmid (Newman Δ*spa*/*sbi* + p*spa*). Correct overexpression of SpA on the *S. aureus* surface was validated by anti-SpA IgY antibodies and flow cytometry (**Fig. 3A**). Moreover, SpA functionality on the bacterial surface was confirmed by studying that an IgG1 isotype control (anti-2,4-dinitrophenol (DNP) antibody) can bind cell-surface-SpA-expressing strains (Newman WT and Newman Δ*spa*/*sbi* + p*spa*), but not Newman Δ*spa*/*sbi* (**Fig. S2A**). As anticipated, anti-DNP IgG3 antibodies did not bind any of the Newman strains (**Fig. S2B**). Next, we compared phagocytosis of these three strains in the presence of monoclonal IgGs directed against WTA. After IgG1 and IgG3 were confirmed to similarly bind to the three Newman strains (**Fig. S2C, D**), we showed that IgG1-mediated phagocytosis was lower for the cell-surface-SpA-expressing strains than for the knockout strain (**Fig. 3B**). Notably, the inhibitory effect of SpA on phagocytosis was even more prominent when SpA was overexpressed (**Fig. 3B**). All three bacterial strains were efficiently phagocytized when labeled with anti-WTA IgG3 antibodies (**Fig. 3C**). Overall, these data suggest that, similar to soluble SpA, also cell-surface SpA reduce IgG-mediated phagocytosis of *S. aureus* by binding to the IgG-Fc region.

**Figure 3.**
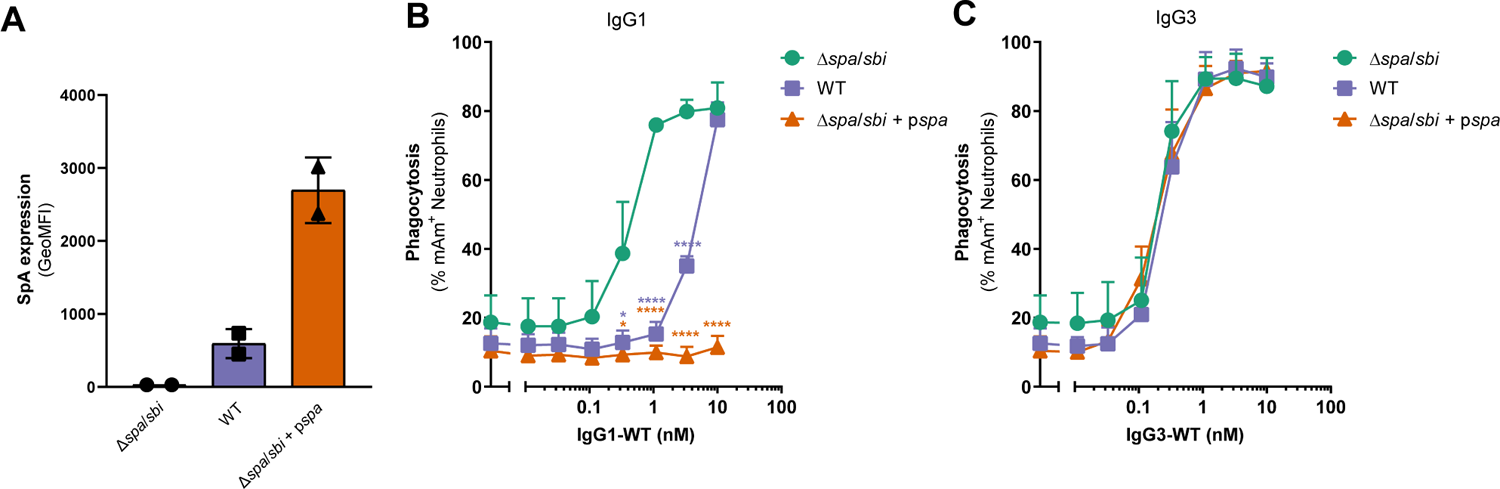
Cell-anchored SpA blocks IgG-mediated phagocytosis of *S. aureus* by binding to IgG-Fc domains. (A) SpA expression on the surface of Newman Δ*spa*/*sbi,* Newman WT, and Newman Δ*spa*/*sbi* + p*spa*, detected with biotinylated-anti-SpA IgY, by flow cytometry; (B, C) Phagocytosis of *S. aureus* Newman Δ*spa*/*sbi*, Newman WT and Newman Δ*spa*/*sbi* + p*spa* strains after incubation of bacteria with a concentration range of anti-WTA IgG1 (B), or IgG3 (C), measured by flow-cytometry. Data are presented as geometric mean fluorescence intensity (GeoMFI) ± SD of two independent experiments (A), or as % of mAm^+^ PMNs or FITC^+^ PMNs ± SD of three independent experiments (B, C). (B, C) Statistical analysis was performed using one-way ANOVA to compare Newman Δ*spa*/*sbi* conditions with Newman WT and Newman Δ*spa*/*sbi* + p*spa* conditions and displayed only when significant as *P ≤ 0.05; ****P ≤ 0.0001.

### Soluble, multi-domain SpA affects binding of bacterium-bound IgG1 to FcγRIIa and FcγRIIIb, but not to FcγRI

Since FcγRs are generally believed to be the main drivers of IgG-mediated phagocytosis, we studied whether SpA could interfere with IgG-FcγRs interactions. While neutrophils mainly express FcγRIIa and FcγRIIIb on their surface, FcγRI is found at low abundance (43, 44). Thus, we focus on the effect of SpA on IgG binding to these three FcγR classes. We performed binding assays of IgG1-labeled Newman Δ*spa*/*sbi* to membrane-bound FcγRs using Chinese hamster ovary (CHO) cell lines that stably express single human FcγRs (37). These include FcγRI, FcγRIIa, and FcγRIIIb and the respective polymorphic variants (FcγRIIa H131, FcγRIIa R131, FcγRIIIb NA1 and FcγRIIIb NA2). We observed that the presence of the SpA constructs did not affect binding of IgG1-coated bacteria to hFcγRI-expressing CHO cells (**Fig. 4A** and **S3A**). However, the multi-domain SpA proteins reduced the binding of IgG1-labeled bacteria to hFcγRIIa-expressing CHO cells (**Fig. 4B, C** and **S3B, C**) and to hFcγRIIIb-expressing CHO cells (**Fig. 4D, E** and **S3D, E**). Curiously, the single B domain also decreased the binding of IgG1-coated bacteria to hFcγRIIIb-expressing CHO cells, although less efficiently than multi-domain SpA (**Fig. 4D, E** and **S3D, E**). Of note, we confirmed that IgG1-bound bacteria were unable to bind untransfected CHO cells (**Fig. S3F, G**). Taken together, these results show that soluble SpA composed of five domains interferes with the binding of IgG1-coated bacteria to membrane bound FcγRIIa and FcγRIIIb, but not with FcγRI, and that SpA-B can only interfere with IgG-FcγRIIIb binding.

**Figure 4.**
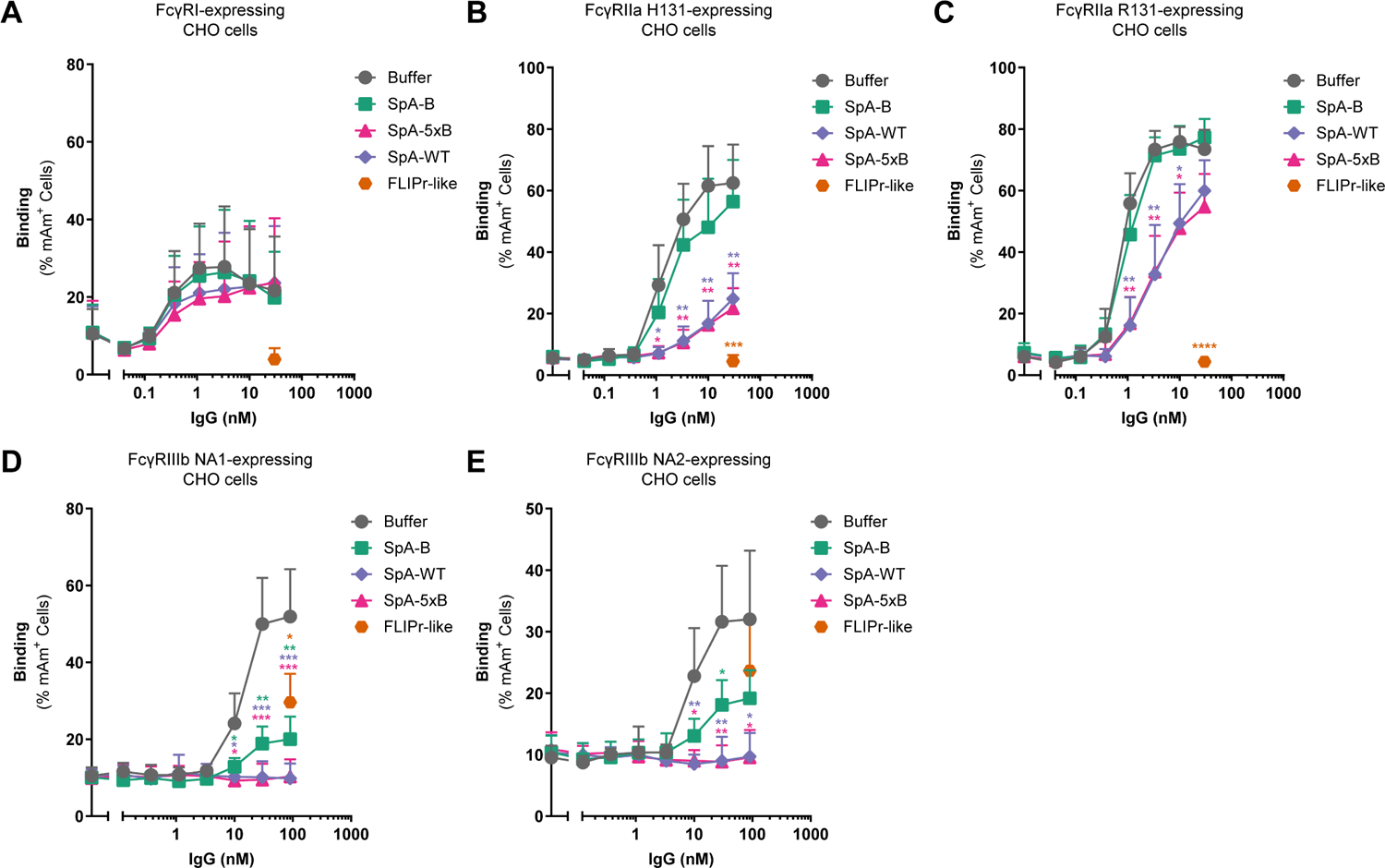
Soluble multi-domain SpA inhibits binding of FcγRIIa and FcγRIIIb to target-bound IgG1. (A-E) Binding of anti-WTA IgG1-labeled *S. aureus* Newman Δ*spa*/*sbi* to hFcγRI-(A), hFcγRIIa H131-(B), hFcγRIIa R131-(C), hFcγRIIIb NA1-(D) and to hFcγRIIIb NA2-expressing CHO cells (E) in absence (buffer; grey) or presence of 200 nM of SpA-B (green), SpA-WT (blue), SpA-5xB (pink) or FLIPr-like (orange), detected by flow-cytometry. Bacteria were washed after incubation with IgG1 to remove unbound antibodies and only after buffer, SpA or FLIPr-like was added. Data are presented as % of mAm^+^ CHO cells ± SD of at least three independent experiments. Statistical analysis was performed using one-way ANOVA to compare buffer condition with SpA-B, SpA-WT, SpA-5xB and FLIPr-like conditions and displayed only when significant as *P ≤ 0.05; **P ≤ 0.01; ***P ≤ 0.001.

### Soluble, multi-domain SpA affects binding of soluble IgG1 to all FcγR classes, except FcγRI, when SpA is in excess

While our assays with target-bound IgG1 indicate that multi-domain SpA blocks FcγRIIa-IgG1 and FcγRIIIb-IgG1 interactions, a previous study showed that SpA does not compete with FcγRIIa to bind soluble IgG1 (26). Although it is more relevant to investigate how SpA competes with FcγR for binding to target-bound IgG than to soluble IgG, we also performed surface plasmon resonance (SPR) experiments to assess the effects of our defined SpA fragments on binding of soluble IgG1 to all FcγR classes and polymorphic variants. FcγRs were coupled to streptavidin biosensors and soluble IgG1 in presence or absence SpA were subsequently injected, at two different IgG:SpA molar ratios (1:1 and 1:5). Contrasting with what was previously reported (26), we found that the multi-domain SpA proteins reduced the interaction of IgG1 to all FcγRs coated on the chip, except to FcγRI, when 1:5 IgG:SpA molar ratio was used (**Fig. S4A**). SpA-B also did not decrease IgG-FcγR interaction (**Fig. S4A**). When instead of 1:5, we used a IgG:SpA molar ratio of 1:1, the presence of SpA altered the IgG-FcγRs binding kinetics, but did not reduce the interaction of IgG1 to any of the FcγRs spotted on the chip (**Fig. S4B**). In fact, under these conditions we found that the multi-domain SpA constructs decreased the on-rates but also the off-rates and hence enhanced the stability of the IgG-FcγR binding (**Fig. S4B**). Altogether, these results show that multi-domain SpA can still interfere with binding of soluble IgG1 to low-affinity FcγRs when SpA is in excess in relation to soluble IgG, but not when IgG and SpA are added at an equimolar ratio.

### Soluble SpA inhibits IgG1-FcRn interactions

Besides extracellular FcγRs, neutrophils also express FcRn inside granular structures (2) (see **Fig. 1A**). The presence of FcRn in neutrophils was shown to be important for efficient IgG-mediated phagocytosis of pneumococci (2). We previously suggested that, upon binding of target-bound IgG to FcγRs on the surface of neutrophils, FcRn is translocated to nascent phagosomes where the low pH promotes binding of FcRn to IgG and facilitates internalization of IgG-opsonized targets (2), additionally, it seems to further promote inflammation in autoimmunity (45). Since SpA and FcRn have an overlapping binding site on the IgG Fc-region (see **Fig. 1B**) and that an analog of the B domain of SpA (the Z domain) was shown to inhibit the binding of FcRn to soluble IgG (46), we also measured the impact of SpA on IgG-FcRn interactions.

We performed flow cytometry experiments in which bacterium-bound IgG1 alone or in combination with each of the SpA constructs, or FLIPr-like, were incubated with FcRn-coated beads at pH 6.0. As expected, all SpA variants reduced IgG1-FcRn interactions, although multi-domain SpA proteins were more effective than SpA-B (**Fig. 5A, B**).

**Figure 5.**
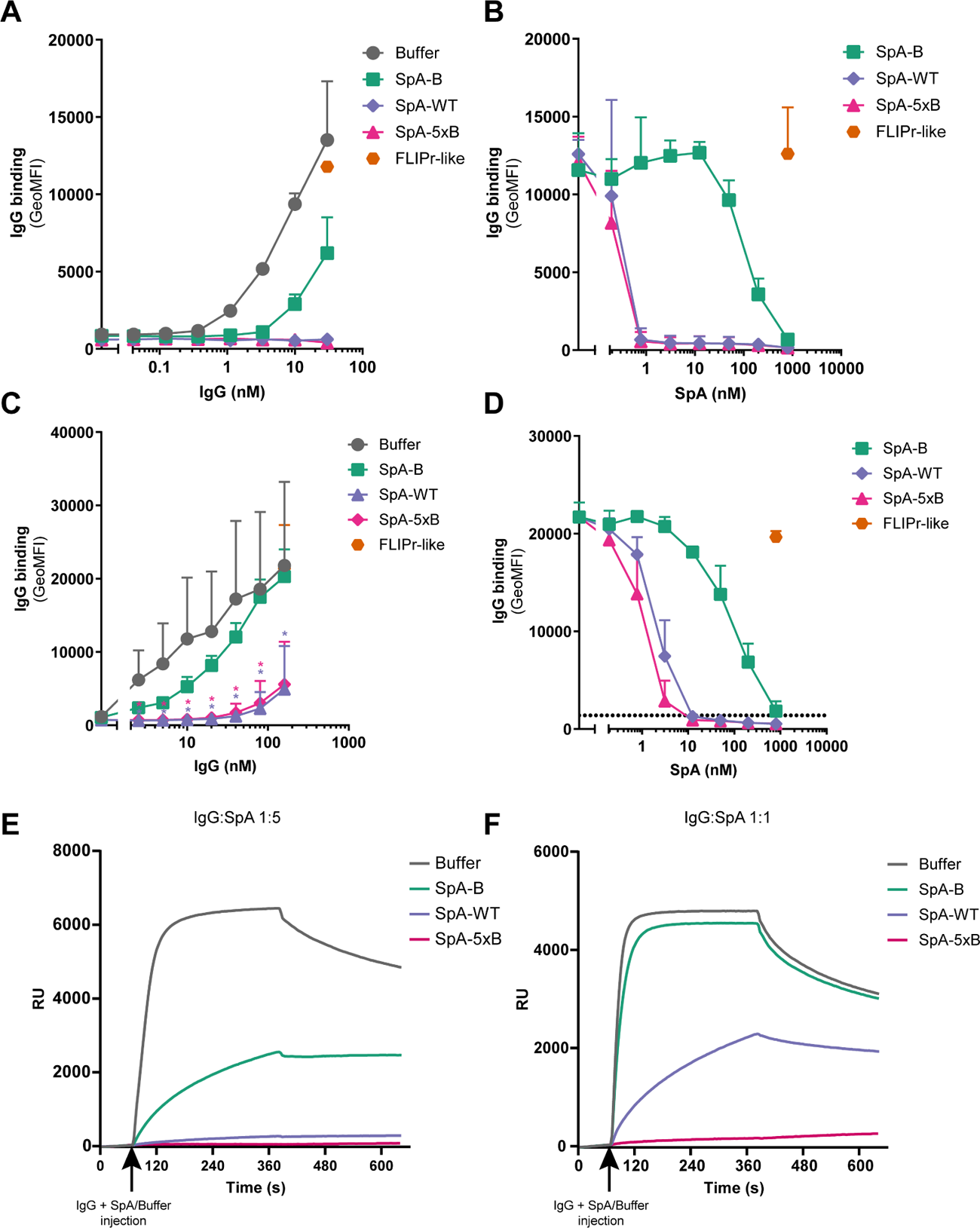
Soluble SpA inhibits binding of FcRn to target-bound and soluble IgG1. (A) Binding of anti-WTA IgG1-labeled *S. aureus* Newman Δ*spa*/*sbi* to FcRn-coated beads at pH 6.0 in absence (buffer; grey) or presence of 200 nM of SpA-B (green), SpA-WT (blue) or SpA-5xB (pink), detected with Alexa Fluor^647^-conjugated goat F(ab’)_2_ anti-human kappa by flow-cytometry. (B) Binding of anti-WTA IgG1-labeled *S. aureus* Newman Δ*spa*/*sbi* bound to FcRn-coated beads in presence of a concentration range of SpA-B (green), SpA-WT (blue) or SpA-5xB (pink), using 10 nM IgG1, detected with Alexa Fluor^647^-conjugated goat F(ab’)_2_ anti-human kappa by flow-cytometry. (C) Binding of a concentration range of IgG1 to FcRn-coated beads in absence (buffer; grey) or presence of 200 nM SpA-B (green), SpA-WT (blue), SpA-5xB (pink) or FLIPr-like (orange), detected with Alexa Fluor^647^-conjugated goat F(ab’)₂ anti-human kappa by flow-cytometry. Statistical analysis was performed using one-way ANOVA to compare buffer condition with SpA-B, SpA-WT, SpA-5xB and FLIPr-like conditions and displayed only when significant as *P ≤ 0.05. (D) Binding of 10 nM of IgG1 to FcRn-coated beads in presence of a concentration range of SpA-B (green), SpA-WT (blue), SpA-5xB (pink) or FLIPr-like (orange), detected with Alexa Fluor^647^-conjugated goat F(ab’)₂ anti-human kappa by flow-cytometry. (E, F) Sensorgram of SPR measurement for binding of 200 nM of IgG1 to FcRn in absence (buffer; grey) or presence of 1 µM (E) or 200 nM (F) of SpA-B (green), SpA-WT (blue) or SpA-5xB (pink). FcRn was first spotted on the sensor and after IgG1 alone or in combination with SpA-B, SpA-WT or SpA-5xB was injected at pH 6.0. Data are presented as mean ± SD of two (A, B, D) or three (C) independent experiments or as response units (RU) of a representative experiment of two independent experiments (E, F).

We also tested whether there was a difference on the effect of SpA on IgG-FcRn binding when IgGs were in solution. We measured binding of soluble IgG1 to FcRn-coated beads in presence or absence of SpA and showed that all SpA variants reduced IgG1-FcRn interactions, although the single domain was less effective than multi-domain SpA proteins (**Fig. 5C, D**). Flow cytometry measurements were corroborated by SPR experiments where biotinylated-FcRn was coupled to streptavidin sensors and soluble IgG1 alone or in combination with each of the SpA proteins was subsequently injected, using a IgG:SpA molar ratio of 1:5 (**Fig. 5E**). However, when IgG1 and SpA were injected at an equimolar ratio, SpA-B lost its ability to block IgG-FcRn interactions (**Fig. 5F**). Overall, these data indicate that SpA directly competes with FcRn for binding IgG and that SpA needs to bind to both Fc-binding sites of an IgG to prevent IgG-FcRn binding. Moreover, these results help to clarify how SpA blocks IgG-mediated phagocytosis.

### Soluble and surface-bound SpA affect phagocytosis of *S. aureus* mediated by naturally occurring antibodies

Finally, we studied the effect of SpA on IgG-mediated phagocytosis in normal human serum (NHS) which contains naturally occurring antibodies against *S. aureus*. The serum was heat-inactivated (HI-NHS) to prevent complement activation. Although multi-domain SpA proteins were more effective than SpA-B, all SpA constructs reduced antibody-mediated phagocytosis (**Fig. 6A, B**), even though HI-NHS comprises many different antibodies, including antibodies that bind SpA at different regions (Fc and/or Fab domains), and also antibodies that do not bind SpA (as IgG3 and IgM from non VH3-type family). We also assessed the impact of cell-surface SpA in reducing phagocytosis in presence of HI-NHS. The efficiency of phagocytosis was reduced when neutrophils were challenged to engulf cell-surface-SpA-expressing strains, when compared with Newman Δ*spa*/*sbi,* in particular when the SpA overexpressing Newman Δ*spa*/*sba+*p*spa* strain was used that resisted IgG-mediated phagocytosis (**Fig. 6C**). Altogether, these data show that SpA can block phagocytosis mediated by naturally occurring antibodies and that the single SpA-B domain is sufficient to affect it.

**Figure 6.**
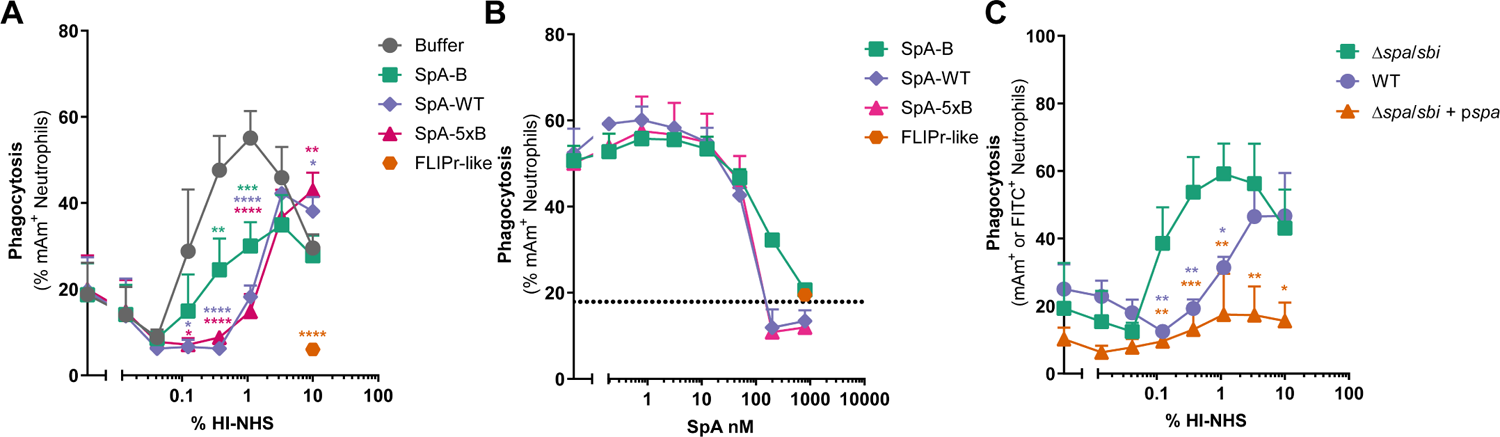
Soluble SpA blocks phagocytosis of *S. aureus* mediated by naturally occurring antibodies. (A) Phagocytosis of *S. aureus* Newman Δ*spa*/*sbi* after incubation of bacteria with a concentration range of heat inactivated human normal serum (HI-NHS), in absence (buffer; grey) or presence of 200 nM soluble SpA-B (green), SpA-WT (blue), SpA-5xB (pink) or FLIPr-like (orange), measured by flow-cytometry. Bacteria, IgG and Buffer/SpA were incubated at the same step. Statistical analysis was performed using one-way ANOVA to compare buffer condition with SpA-B, SpA-WT, SpA-5xB and FLIPr-like conditions and displayed only when significant as *P ≤ 0.05; **P ≤ 0.01; ***P ≤ 0.001; ****P ≤ 0.0001. (B) Phagocytosis of *S. aureus* Newman Δ*spa*/*sbi* after incubation of bacteria with 1% HI-NHS in presence of a concentration range of SpA-B (green), SpA-WT (blue) or SpA-5xB (pink), measured by flow-cytometry. Bacteria, IgG and Buffer/SpA were incubated at the same step. The black dotted line shows the background fluorescence from bacteria that were not incubated with IgG. (C) Phagocytosis of *S. aureus* Newman Δ*spa*/*sbi*, Newman WT and Newman Δ*spa*/*sbi* + p*spa* strains after incubation of bacteria with a concentration range of HI-NHS, measured by flow-cytometry. Statistical analysis was performed using one-way ANOVA to compare Newman Δ*spa*/*sbi* conditions with Newman WT and Newman Δ*spa*/*sbi* + p*spa* conditions and displayed only when significant as *P ≤ 0.05; **P ≤ 0.01; ***P ≤ 0.001. Data are presented as % of mAm^+^ PMNs ± SD or FITC^+^ PMNs ± SD of three (A, C) or two (B) independent experiments.

## Discussion

Antibodies can help to resolve infections by inducing Fc-effector functions after binding to bacterial surfaces (1). The Fc domains of IgG-labeled bacteria are recognized by FcγRs that are expressed on the surface of innate immune cells, e.g. neutrophils, which engulf and kill bacteria intracellularly. Next to extracellular FcγRs, neutrophils also express FcRn intracellularly, which showed to facilitate IgG-mediated phagocytosis (2). In this study, we made two important discoveries that help to understand how SpA from *S. aureus* blocks IgG-mediated phagocytosis: first, we revealed that SpA interferes with the binding of IgG to FcγRIIa and FcγRIIIb; second, we found that SpA blocks the interaction between IgG and FcRn. Our findings contribute for a better understanding of the immune evasion mechanisms of *S. aureus*. Moreover, our study supports that FcRn, besides FcγRs, also has an important role in phagocytosis.

This work confirms that both soluble and cell-attached SpA efficiently block FcR-mediated phagocytosis of *S. aureus* by human neutrophils and, more importantly, it shows that this is because SpA blocks the binding of FcγRIIa, FcγRIIIb and FcRn to target-bound IgGs. Although SpA has been known to block phagocytosis for a long time (23), the molecular mechanism behind it was not clarified. While SpA was shown to block binding of IgG-labeled surfaces to Fc receptor-expressing cells (23, 25, 47), soluble murine FcγRI and human FcγRIIa were demonstrated not to compete with SpA for binding to IgG (26). Here, we confirm that SpA does not affect the binding of the high-affinity human FcγRI to IgG1. However, we show that SpA decreases the binding of the low-affinity receptors FcγRIIa and FcγRIIIb to IgG1. These conflicting results might be explained by the fact that, instead of soluble FcγRs, we use surface-bound FcγRs as they better resemble membrane FcγRs. Soluble FcγRs are likely less constrained in their mobility, which may facilitate their binding to SpA-bound IgG molecules.

It has been presumed that SpA, by binding antibodies, would simply sequester their Fc sites and, thus, preclude Fc recognition by phagocytic cells (23, 47, 48). Here, we clarify that the binding of SpA to IgG1 likely causes a suboptimal sterical conformation that affects the binding of low-affinity FcγRs but not high-affinity FcγRs. Importantly, we show that SpA also prevents FcRn from binding IgG. Contrarily to FcγRs, that bind IgG-Fc in a structurally distant site from the SpA binding site, FcRn interacts with IgG at the exact same site as SpA. Thus, while SpA-IgG interactions likely prevent binding of FcγRs due to steric hindrance, FcRn should compete directly with SpA for binding IgG. Although an analog of the B domain of SpA (the Z domain) was previously shown to inhibit the binding of FcRn to soluble IgG (46), here we associate the effect of SpA on blocking IgG-FcRn interactions with its anti-phagocytic properties. Importantly, by blocking binding of FcRn to IgG, SpA may also interfere with other important functions of FcRn. In addition to its role in IgG-phagocytosis by neutrophils (2), FcRn also mediates the transfer of IgG from the mother to her fetus (49) and extends the serum half-life of IgG (5, 50). More recently, FcRn was also found to regulate antigen presentation (51), antigen cross-presentation (52, 53) and secretion of cytotoxicity-promoting cytokines by dendritic cells (52). Therefore, we expect that SpA has a much broader immunomodulatory action than initially anticipated.

This study also provides a rationale for the multiplicity of repeating Ig-binding domains of SpA produced by *S. aureus*. In fact, we show that SpA needs to be composed of multiple Ig-binding domains to efficiently block IgG1-mediated phagocytosis and to decrease the binding of IgG1 to FcγRIIa-coated surfaces. It is possible that a multi-domain SpA molecule that is bound to IgG1-opsonized bacteria can still bind to soluble IgGs, forming IgG-SpA complexes that make IgG-Fc tails inaccessible to FcγRs. However, experiments where the bacteria were first incubated with IgGs and then washed suggest that SpA binds to bacterium-bound IgGs to block IgG-FcγRs interactions and, consequently, phagocytosis. Thus, a more plausible hypothesis is that SpA needs multiple IgG-binding domains to cause steric hindrance and mask the FcγR binding site on IgG1-opsonized bacteria. This hypothesis is supported by the fact that SpA composed of five domains binds IgG with a 1:1 stoichiometry (21), which suggests that two of the five domains of SpA bind to both sites of IgG-Fc, leaving the other three domains free to cover the region in IgG where FcγRs bind. However, this theory does not explain why the single SpA-B domain affects IgG-FcγRIIIb binding. We speculate that this is a consequence of a slight change to the CH2 configuration of the IgG that is induced by the binding of SpA-B, which has a particularly strong impact (relative) on the lowest-affinity FcγR.

Although further studies are needed to clarify how SpA-B affects IgG2- and IgG4-mediated phagocytosis and whether the target of the antibody influences FcγR and/or FcRn recognition, we suggest that SpA-B may be used as a research tool to assess which FcR(s) drive IgG-mediated phagocytosis. Because the effect of SpA-B on phagocytosis mediated by anti-WTA IgG1 was very minor, it is likely that this antibody mediates phagocytosis mainly by triggering FcγRIIa. Conversely, since the single SpA-B domain was sufficient to effectively reduce phagocytosis mediated by naturally occurring antibodies, FcRn and/or FcγRIIIb may play a more essential role in the phagocytosis mediated by other antibody types.

Around 85% of SpA produced by *S. aureus* is anchored to the cell wall of the bacteria (14). We envision that cell attached SpA might inhibit IgG-mediated phagocytosis of *S. aureus* by the same mechanisms described here for soluble SpA. However, we also speculate that cell attached SpA could induce an additional inhibitory mechanism by covering *S. aureus* surface with antibodies, creating a shield that prevents anti-*S. aureus* antibodies from reaching the bacterial surface and/or that masks their binding sites, as suggested before (23). Another hypothesis is that their binding sites are already occupied by antibodies that simultaneously bind to cell-surface SpA (via Fc-domain) and to their target antigen on the bacterial surface (via Fab-domain), the so-called phenomenon “bipolar bridging”.

Our study also provides a rational for the design of therapeutic antibodies against staphylococcal infections. In line with our previous study (21), we show here that IgG3 antibodies are unaffected by the presence of SpA, and thus are more potent to mediate phagocytosis and killing of *S. aureus* by neutrophils than IgG1. Therefore, we suggest that monoclonal antibodies against *S. aureus* surface should be developed as IgG3 antibodies. The fact that IgG3 antibodies are the most effective IgG subclass to trigger immune effector functions also supports this selection.

In conclusion, this study unveils how SpA blocks IgG-mediated phagocytosis, which improves our understanding of the immune evasion strategies of *S. aureus* and may help the development of therapeutic options to tackle staphylococcal infections.

## Acknowledgments

We thank Dr. Annette M. Stemerding for fruitful discussions.

