## Supplemental Data for "Staphylococcal protein A inhibits IgG-mediated phagocytosis by blocking the interaction of IgGs with FcγRs and FcRn"

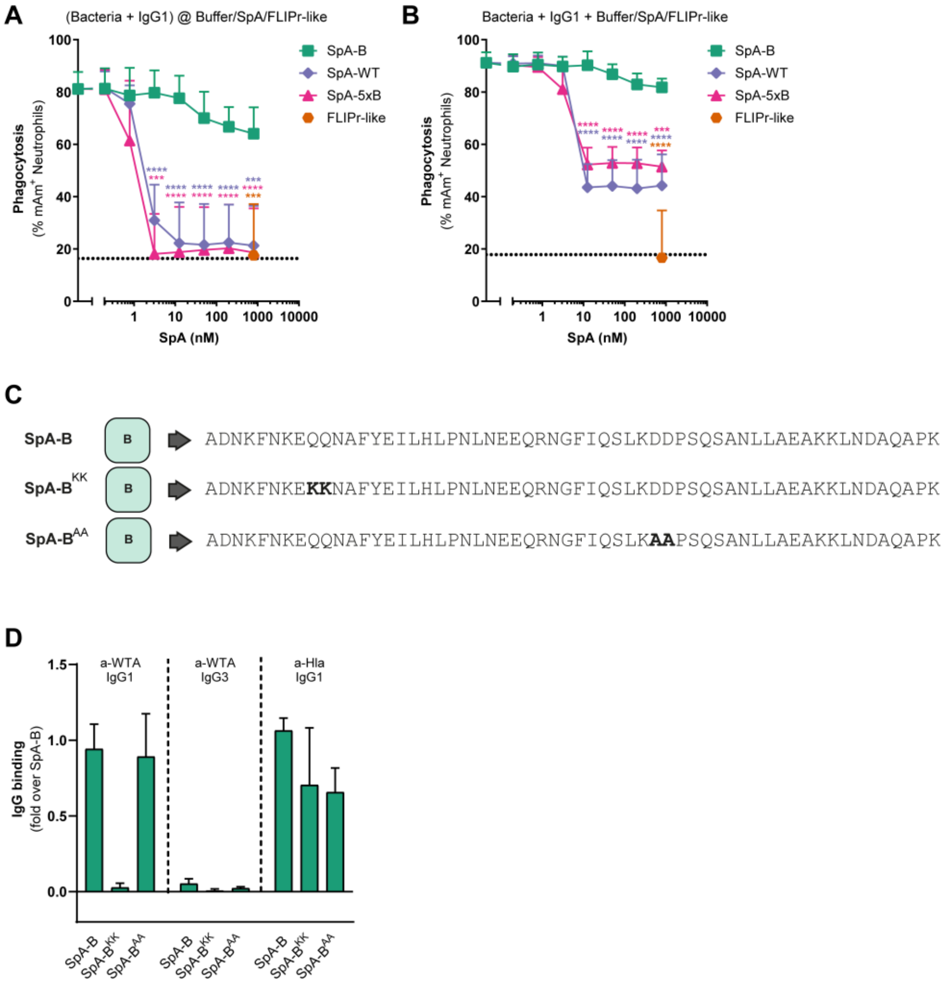


**Figure S1. Soluble multi-domain SpA blocks antibody-mediated phagocytosis of *S. aureus* by binding to IgG via Fc domain.** (A) Phagocytosis of *S. aureus* Newman Δ*spa*/*sbi* after incubation of bacteria with 10 nM anti-WTA IgG1, followed by incubation with a concentration range of SpA-B (green), SpA-WT (blue), SpA-5xB (pink) or FLIPr-like (orange), measured by flow-cytometry. Bacteria were washed after incubation with IgGs to remove unbound IgG and only after buffer or SpA was added. (B) Phagocytosis of *S. aureus* Newman Δ*spa*/*sbi* after incubation of bacteria with 10 nM anti-WTA IgG1, in presence of a concentration range of SpA-B (green), SpA-WT (blue), SpA-5xB (pink) or FLIPr-like (orange), measured by flow-cytometry. Bacteria, IgG and buffer/SpA were incubated at the same step. The black dotted lines show the background fluorescence from bacteria that were not incubated with IgG. (C)

Schematic representation of the recombinant SpA-B proteins used in this experiment. SpA-B binds to antibody Fc and Fab regions while SpA-$B^{KK}$ only binds to IgG-Fab region and SpA-$B^{AA}$ only binds IgG-Fc region. The amino acid sequences of the three protein are indicated and the KK and AA mutations are highlighted in bold. (D) Binding of 1 µg/mL anti-WTA IgG1, anti-WTA IgG3 and anti-Hla IgG1 to SpA-B (wild-type), SpA-$B^{KK}$ (only binds to IgG-Fab region; Q9K and Q10K mutations), or SpA-$B^{AA}$ (only binds to IgG-Fc region; D36A and D37A mutations), coated on a ELISA plate. IgG binding was detected with HRP-labeled goat $F(ab')_2$ anti-human kappa. Data are presented as mean ± SD of at least three independent experiments. (A, B) Statistical analysis was performed using one-way ANOVA to compare buffer condition with SpA-B, SpA-WT, SpA-5xB and FLIPr-like conditions and displayed only when significant as ***P ≤ 0.001; ****P ≤ 0.0001.

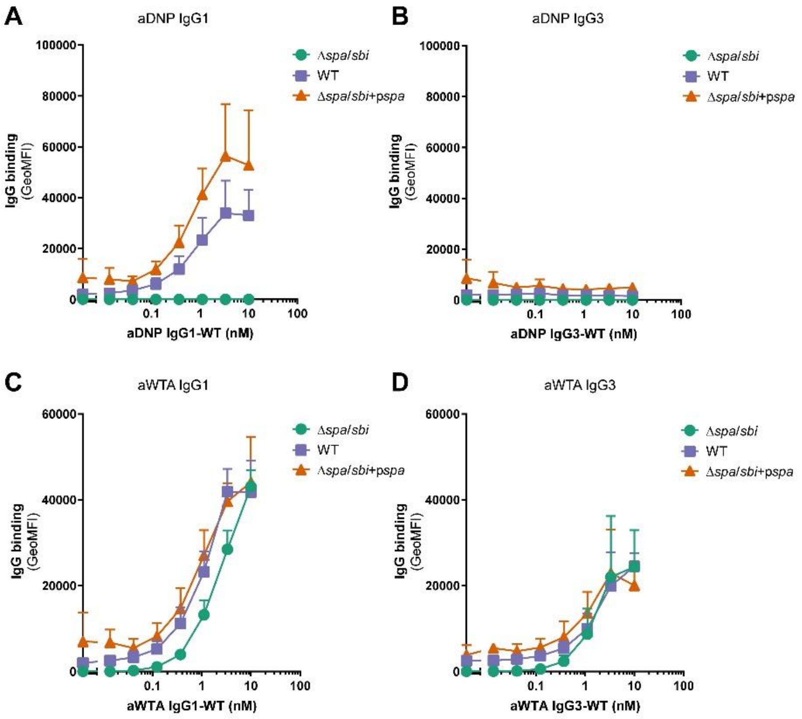


**Figure S2. Characterization of Newman strains.** (A-D) Binding of a concentration range of anti-DNP IgG1 (A), anti-DNP IgG3 (B), anti-WTA IgG1 (C), or anti-WTA IgG3 (D) to Newman Δ*spa*/*sbi*, Newman WT and Newman Δ*spa*/*sbi* + p*spa* strains, detected with Alexa Fluor$^{647}$-conjugated goat F(ab')$_2$ anti-human kappa by flow-cytometry. Data are presented as geometric mean fluorescence intensity (GeoMFI) ± SD of at least two independent experiments.

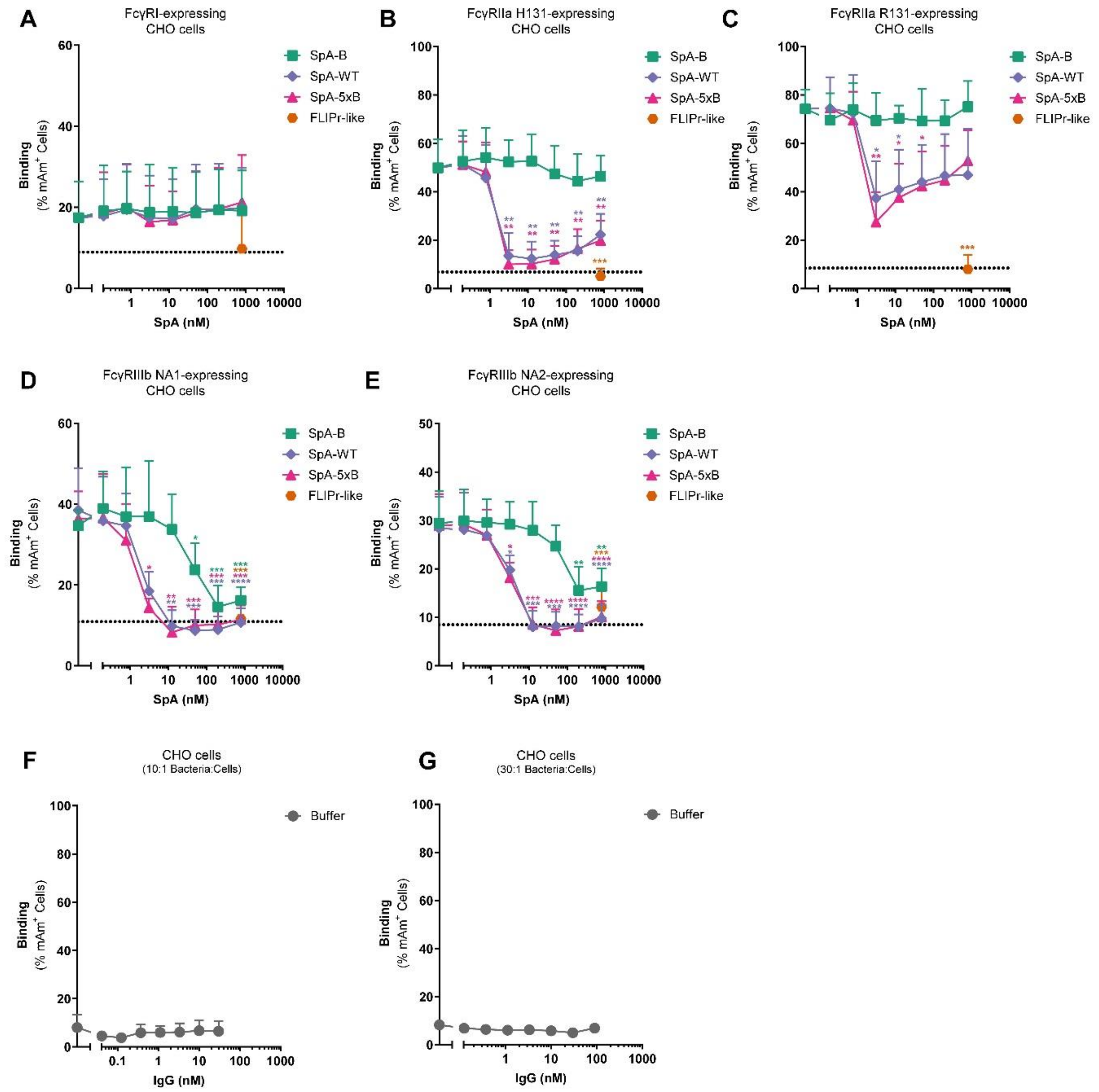


**Figure S3. Soluble multi-domain SpA inhibits binding of FcγRIIa and FcγRIIIb to target-bound IgG1.** (A-E) Binding of anti-WTA IgG1-labeled *S. aureus* Newman Δ*spa*/*sbi* to hFcγRI- (A), hFcγRIIa H131- (B), hFcγRIIa R131- (C), hFcγRIIIb NA1- (D) and to hFcγRIIIb NA2-expressing CHO cells (E) in presence of a concentration range of SpA-B (green), SpA-WT (blue), SpA-5xB (pink) or FLIPr-like (orange), detected by flow-cytometry. Bacteria were washed after incubation with 10 nM IgG1 (A-C) or 30 nM IgG1 (D, E) to remove unbound antibodies and only after buffer, SpA or FLIPr-like was added. (F, G) Binding of anti-WTA IgG1-labeled *S. aureus* Newman Δ*spa*/*sbi* to untransfected CHO cells, when bacteria:cell ratio is of 10:1 (F) and of 30:1 (G), detected by flow-cytometry. Data are presented as % of mAm$^{+}$

CHO cells ± SD of at least three independent experiments. (A-E) Statistical analysis was performed using one-way ANOVA to compare buffer condition with SpA-B, SpA-WT, SpA-5xB and FLIPr-like conditions and displayed only when significant as *$P \leq 0.05$; **$P \leq 0.01$; ***$P \leq 0.001$; ****$P \leq 0.0001$.

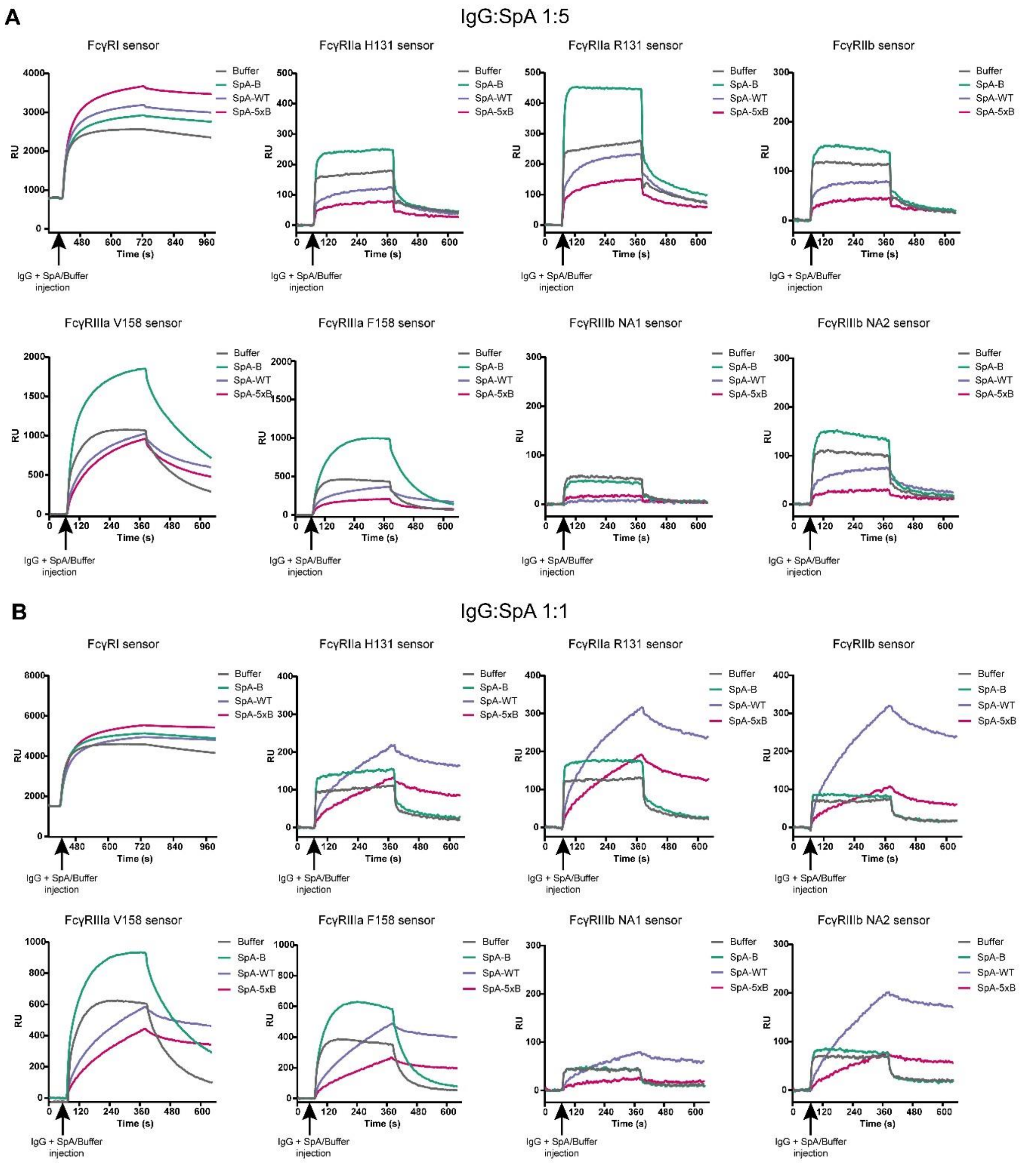


**Figure S4. Soluble multi-domain SpA reduces binding of all FcγRs, expect FcγRI, to soluble IgG1 when SpA is in excess.** (A, B) Sensorgram of SPR measurement for the binding of 200 nM IgG1 to FcγRI, FcγRIIa H131, FcγRIIa R131, FcγRIIb, FcγRIIIa V158, FcγRIIIa F158, FcγRIIIb NA1 and FcγRIIIb NA2 in absence (grey) or presence of 1 µM (A) or 200 nM (B) of SpA-B (green), SpA-WT (blue) or SpA-5xB (pink). FcγRs were first spotted on the sensor and after IgG1 alone or in combination with SpA-B, SpA-WT or SpA-5xB was injected.

Data are presented as response units (RU) of a single representative experiment of two independent experiments.
